# Microglial refinement of A-fibre projections in the postnatal spinal cord dorsal horn is required for normal maturation of dynamic touch

**DOI:** 10.1101/2021.03.22.436389

**Authors:** Yajing Xu, Stephanie C. Koch, Alexander Chamessian, Qianru He, Mayya Sundukova, Paul Heppenstall, RuRong Ji, Maria Fitzgerald, Simon Beggs

## Abstract

Sensory systems are shaped in postnatal life by the refinement of synaptic connections. In the dorsal horn of the spinal cord, sensory circuits undergo postnatal activity dependent reorganisation, including the retraction of primary afferent A-fibres from superficial to deeper laminae which is accompanied by decreases in cutaneous sensitivity. Here we show that microglia, the resident immune cells in the CNS, phagocytose A-fibre terminals in superficial laminae in the first weeks of life. Genetic perturbation of microglial engulfment at that time prevents the normal process of A-fibre retraction, resulting in increased sensitivity of dorsal horn cells to dynamic tactile cutaneous stimulation, and behavioural hypersensitivity to dynamic touch. Thus, functional microglia are necessary for normal postnatal development of dorsal horn sensory circuits. In the absence of microglial engulfment, superfluous A-fibre projections remain in the dorsal horn and the balance of sensory connectivity is disrupted, leading to lifelong hypersensitivity to dynamic touch.

**Impact statement:** Microglia phagocytose superfluous A-fibres in the superficial spinal dorsal horn during normal development, the disruption of which leads to long term aberrant dynamic touch processing and behaviour.

## Introduction

The neonatal spinal dorsal horn differs substantially from that in adults and undergoes extensive structural and functional reorganisation over the postnatal period. One notable change is the termination zone of primary afferent A-fibres, the large myelinated afferents that encompass many low threshold cutaneous tactile afferents ^1^. These afferent terminals occupy both superficial and deep laminae of the dorsal horn in neonatal rodents and gradually retract to terminate in deeper laminae III-IV by the end of the 3rd postnatal week and in adulthood ^2–4^. This retraction of A-fibre terminals is accompanied by a reduction in dorsal horn cell receptive field sizes on the skin, as well as a decline in tactile sensitivity and an increase in reflex behaviour precision, with similar changes in somatosensory behaviour observed in human infants (Fitzgerald 1985, 2015; Fitzgerald et al. 1988). This suggests that structural refinement in the dorsal horn likely underlies behavioural maturation.

The process of A-fibre terminal retraction in the first weeks of life is activity dependent: Blocking neuronal input through spinal NMDAR inhibitors or increasing the noise of neuronal input through random vibration to the skin over extended period both prevented the normal retraction of A-fibres ^8,9^. However, the exact mechanism underlying the retraction of A-fibre terminals is not known. Microglia cells, the major phagocytes in the CNS, have been shown to remove superfluous neurons both by driving apoptosis, removing apoptotic cells, and phagocytosing synapses and neurites during postnatal refinement ^10^. To date, such studies have been largely restricted to the brain ^11–15^, with two studies reporting a role of microglia in the postnatal development of spinal cord ventral horn motor circuits ^16,17^. Whether microglia are also involved in the maturation of somatosensory circuits in the dorsal horn is not known. Here we hypothesise that the retraction of A-fibres from superficial laminae in the postnatal period is driven by microglia which prune A-fibre terminals in the dorsal horn as part of normal postnatal development of spinal sensory circuits.

Microglia undergo postnatal maturation during which they not only change in density and morphology but also alter their transcriptional and functional identity ^18,19^. Brain microglia show particularly high expression of lysosome associated genes at P4/5 suggesting a specialised role of microglial phagocytosis during development ^20^. In addition, microglia exhibit spatial heterogeneity, as they populate different brain regions at different rates postnatally, and express distinct local genetic profiles and phenotype in adulthood ^21–23^. So far, most research has focused on the brain and relatively little is known about spinal cord microglia.

To test whether A-fibre terminals in the developing spinal dorsal horn are pruned by microglial phagocytosis, we used a transgenic mouse line that expresses tdTomato (tdT) under the *Vglut1* promoter which labels a subset of A-fibres ^24^ and mapped morphological changes in dorsal horn microglia across the first two postnatal weeks using immunofluorescence and confocal microscopy. We next tested whether normal microglial phagocytosis was required for the A-fibre pruning. Constitutive knock-out of the gene *Tmem16f* in microglia was shown to reduce microglial phagocytosis and motility in the adult spinal cord in a neuropathic pain model, resulting in reduced pain behaviour ^25^. Therefore, we blocked microglial phagocytosis during the first postnatal week using a tamoxifen inducible Cre-mediated deletion of the *Tmem16f* gene in microglia and measured the effects on A-fibre pruning, the subsequent maturation of dorsal horn synaptic connections, and behavioural reflex sensitivity to mechanical skin stimulation. The results suggest that microglia prune A-fibre terminals in the developing spinal dorsal horn and that this postnatal microglial refinement of A-fibre terminals is required for normal somatosensory maturation.

## Results

### Dorsal horn microglia phagocytose A-fibre terminals during postnatal development

In the developing spinal cord, tactile-encoding A-fibres initially project throughout the dorsoventral extent of the dorsal horn and refine over the first few postnatal weeks to terminate in deeper laminae, segregated from the more superficial terminals of noxious-encoding C-fibres ^8^. We confirmed this using a transgenic reporter mouse in which tdTomato is expressed in vesicular glutamate transporter 1 (*Vglut1*) expressing neurons (VGLUT1-tdT), a subpopulation of large myelinated sensory neurons with features consistent with Aβ-low threshold mechanoreceptors (LTMRs) ^24^. This tdT-expression was sparse at P0 and increased with age, likely due to the increasing developmental expression profile of *Vglut1* ^26,27^ (Figure 1—figure supplement 1a). Despite this, the mice clearly revealed A-fibre terminals with flame-shaped arbors extending into laminae I-II up to P7 (Figure 1a), which are no longer present in these superficial laminae by P28, consistent with previous findings ^8^.

**Figure 1.**
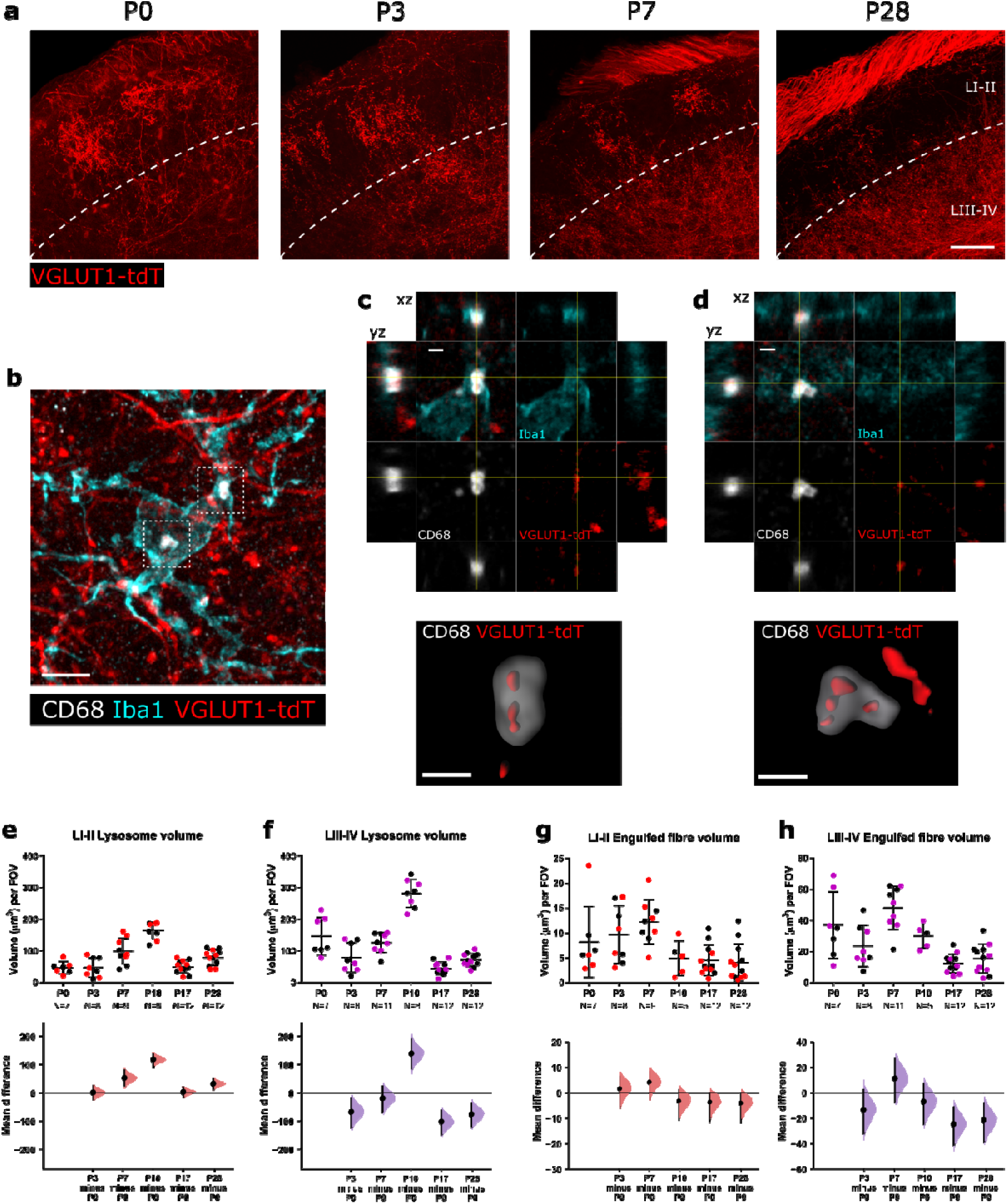
Spinal dorsal horn microglia engulf A-fibres during normal postnatal development. **a**. VGLUT1-tdTomato (red) expression in spinal laminae I-II decreases across age. Dashed white line indicates border between lamina II and lamina III. Scale bar = 50_μ_m. **b**. Representative z-projected super-resolution image of A-fibre engulfment by microglia within the cell body. White inset box show location of higher magnification panels in **c** and **d**. Scale bar = 5 _μ_m. **c, d**. High magnification images of microglial A-fibre engulfment in **b** stained for microglia (Iba1, cyan), microglial lysosomes (CD68, white) and endogenously fluorescent A-fibres (VGLUT1-tdT, red). Cross-hairs show position of the xz and yz side-view panels. Bottom panels show surface rendering of the super-resolution image revealing pieces of tdT labelled fibres engulfed inside lysosome. Scale bar = 1 _μ_m. **e**. Microglial lysosome volume peaks at P10 and decreases thereafter for LI-II. P3 vs. P0: mean difference 0.48 [95% CI 23.52, 24.35]; P7 vs. P0: mean difference 52.19 [95% CI 23.71, 79.47]; P10 vs. P0: mean difference 117.16 [95% CI 92.26, 137.61]; P17 vs. P0: mean difference 2.52 [95% CI - 15.07, 18.38]; P28 vs. P0: mean difference 31.84 [95% CI 11.42, 49.16]. Field of view (FOV) = 245_μ_m x 65_μ_m. **f**. Microglial lysosome volume peaks at P10 and decreases thereafter for LIII-IV. P3 vs. P0: mean difference −66.67 [95% CI −121.07, −18.14]; P7 vs. P0: mean difference −19.48 [95% CI −67.54, 21.66]; P10 vs. P0: mean difference 136.02 [95% CI 82.89, 184.41]; P17 vs. P0: mean difference −100.85 [95% CI −145.40, −62.42]; P28 vs. P0: mean difference −74.95 [95% CI −119.97, −36.61]. Field of view (FOV) = 245_μ_m x 65_μ_m. **g**. Engulfed fibre volume peaks at P7 and decreases thereafter for LI-II. P3 vs. P0: mean difference 1.46 [95% CI −6.86, 6.29]; P7 vs. P0: mean difference 4.05 [95% CI −4.11, 8.18]; P10 vs. P0: mean difference −3.28 [95% CI −10.76, 1.11]; P17 vs. P0: mean difference −3.69 [95% CI −11.57, −0.10]; P28 vs. P0: mean difference −4.11 [95% CI −11.79, −0.38]. Field of view (FOV) = 245_μ_m x 65_μ_m. **h**. Engulfed fibre volume peaks at P7 and decreases thereafter for LIII-IV. P3 vs. P0: mean difference −13.47 [95% CI −31.97, 2.15]; P7 vs. P0: mean difference 11.03 [95% CI −7.37, 26.33]; P10 vs. P0: mean difference −7.05 [95% CI −24.52, 6.62]; P17 vs. P0: mean difference −24.55 [95% CI −40.82, - 11.27]; P28 vs. P0: mean difference −21.49 [95% CI −37.93, −7.92]. Field of view (FOV) = 245_μ_m x 65_μ_m. N-numbers as indicated. Black and colourful data points indicate females and males respectively.

We hypothesised that microglia might be involved in this refinement process by removing A-fibres in the superficial laminae via phagocytosis and determined the phagocytic activity of dorsal horn microglia over this period using the lysosome associated molecular marker CD68 (Figure 1b-f, Figure 1—figure supplement 1i-k). CD68 lysosome volume increased in both laminae I-II and III-IV over the postnatal period, reaching a peak at P10 and declining thereafter at P17 and P28 (LI-II P10 vs P0 unpaired mean difference 117.16 [95.00% CI 92.26, 137.61], LIII-LIV unpaired mean difference 136.02 [95.00% CI 82.89, 184.41]. It is known that the dorsal horn undergoes developmental apoptosis around birth, extending into the first postnatal week. Therefore, to determine whether the rise in CD68 could be contributed to engulfment of apoptotic cells, we also quantified the number of apoptotic cells and phagocytic cups per microglia. In contrast to the rise in CD68, the number of phagocytic cups per microglia decreased over the first postnatal week concomitant with a decrease in apoptotic cell counts (Figure 1—figure supplement 2), suggesting that the increase in CD68 is not due to phagocytosis of apoptotic cells but other material (Phagocytic cups: F(3, 12)=10.97, P=0.0009, apoptotic cells: F(3, 12)=14.18, P=0.0003). In parallel, microglial density and their ratio to neurons increased over this period (Figure 1—figure supplement 1b-h). Together this suggests that the first postnatal week represents a distinct period of microglial activity, characterised by high levels of microglial CD68.

We next asked whether the high level of CD68 expression in microglia during the first postnatal week is associated with the refinement of neural connectivity in the dorsal horn through the removal of superfluous A-fibre terminals. Engulfment of A-fibres was measured by quantifying VGLUT1-tdT fluorescence volume within CD68-positive lysosomes inside Iba1-labelled microglia cells (Figure. 1 b-d, g-h).

Consistent with the postnatal increase in microglial lysosome volume, the volume of engulfed A-fibres was high during the first postnatal week, peaking at P7 and decreasing thereafter in both laminae I-II, (unpaired mean difference P28 vs. P0 - 4.11 [95% CI −11.79, −0.38]) and laminae III-IV (unpaired mean difference P28 vs. P0-21.49 [95% CI −37.93, −7.92]). This effect was more pronounced in the deeper laminae LIII-IV, which has higher volumes of fibre engulfment at all ages compared to laminae I-II likely due to the higher density of A-fibres in the deeper laminae.

### Microglial activity is required for normal A-fibre pruning in laminae I-III

Having established that microglia participate in postnatal dorsal horn remodeling of A-fibre afferent terminals, we sought to determine whether microglial function is necessary for this process to occur normally. The phospholipid scramblase TMEM16F is required for normal microglial phagocytic activity^10^. To address this question we disrupted microglial function postnatally using a Cre-inducible conditional knocking out of *Tmem16f* in microglia cells ^25^, and observed the structural, functional and behavioural consequences in adult animals. A *R26*^*LSL-Ai9*^ reporter line was used to label *Cxc3cr1*^*Cre*^-expressing microglial cells ^28,29^ and *Thy1*^*eGFP*^ allele was used to identify A-fibres ^30,31^.

To control for off-target effects of Cre expression and tamoxifen administration, both cKO mice and control mice (*Cx3cr1*^*CreER/+*^; *Tmem16f*^*fl/fl*^; *R26*^*LSL-Ai9*^; *Thy1*^*eGFP*^ and *Cx3cr1*^*CreER/+*^; *Tmem16f*^*+/+*^; *R26*^*LSL-Ai9*^; *Thy1*^*eGFP*^ respectively) received 4-hydroxytamoxifen (4-HT) daily from P1 to P3 and were assessed at 3-4 months old. Specificity of Cre expression was confirmed with *R26*^*LSL-Ai9*^ tdT expression.

### Adult microglial Tmem16f cKO mice have increased VGLUT1+ terminals in superficial but not deep laminae

Postnatal deletion of microglial *Tmem16f* resulted in adults with increased A-fibre terminal occupancy in the superficial dorsal horn, as revealed by both Thy1-GFP expression and VGLUT1 immunohistochemistry (Figure 2a-b, d). As Thy1-GFP is expressed in only a small number of sensory neurons and varied across animals, all subsequent quantification was performed using VGLUT1 immunolabelling. Synaptic density of this increased input was quantified using VGLUT1 immunohistochemistry in laminae I-III (Figure 2a upper panels). *Tmem16f* cKO mice had greater primary afferent VGLUT1 synaptic density throughout the dorsal horn than control mice as measured by both total synapse volume and synapse number (Figure 2b) (unpaired mean difference VGLUT1 volume: 799.28 [95.00% CI 450.09, 1136.35], VGLUT1 number: 533.67 [95.00% CI 356.12, 695.50]). In contrast, local inhibitory VGAT synapse density was unaltered (Figure. 2a, c) (unpaired mean difference VGAT volume: −168.31 [95.00% CI −562.61, 253.44], VGAT number: −159.19 [95.00% CI 416.42, 148.88]).

**Figure 2.**
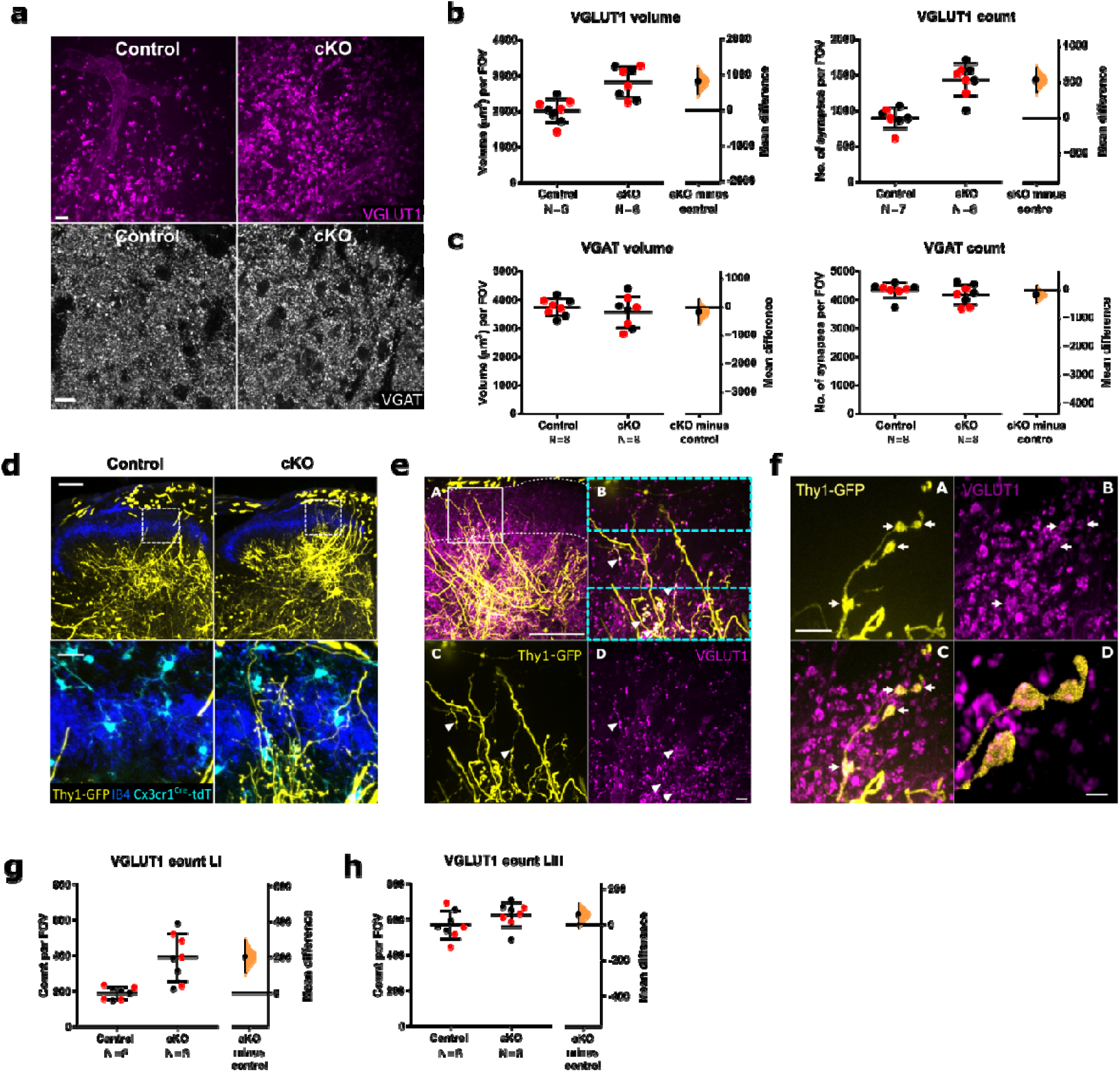
Neonatal *Tmem16f* cKO in microglia increases dorsal horn A-fibre terminals. **a**. Representative images of VGLUT1 and VGAT puncta from the spinal dorsal horn of adult *Tmem16f* control and cKO animals. Field of view (FOV) = 94_μ_m x 94_μ_m (VGLUT1), 96_μ_m x 96_μ_m (VGAT). Scale bars = 10_μ_m. **b**. VGLUT1 puncta volume and count were both increased in adult *Tmem16f* cKO animals compared to controls. Control vs cKO volume: mean difference 799.28 [95.00% CI 450.09, 1136.35]; control vs cKO count: mean difference 533.67 [95.00% CI 356.12, 695.50]. **c**. VGAT puncta volume and count were not significantly different between adult *Tmem16f* cKO animals and controls. Control vs cKO volume: mean difference −168.3105 [95.00% CI −562.61, 253.44]; control vs cKO count: mean difference −159.19 [95.00% CI −416.42, 148.88]. **d**. Thy1-GFP labelled A-fibres (yellow) are present in the superficial dorsal horn (delimited by IB4, blue) of adult cKO mice, but not in adult controls. tdTomato-labelled microglia (cyan) are present in both cKO and control animals. Lower panels show high power images of the boxed areas in the upper panels. Scale bar = 100_μ_m in the top panel and 20_μ_m in the lower panel. **e**. Thy1-GFP labelled A-fibres (yellow) in the superficial dorsal horn laminae of adult *Tmem16f* cKO mice express VGLUT1 (magenta). (A) Low magnification image showing superficially-projecting Thy1-GFP labelled fibres. White box indicates where images were taken for analysis in **a-c**. (B) High magnification of boxed region in A. Cyan boxes indicate the cropped areas used for analysis in **g** and **h**. (C) Thy1-GFP labelling. (D) VGLUT1 immunoreactivity. Scale bar in A = 100µm. Scale bar in D = 10µm. **f**. A-D. High-magnification examples for VGLUT1-expression (magenta) in Thy1-GFP labelled A-fibres (yellow) in adult *Tmem16f* cKO mice. Scale bar in A = 20_μ_m, scale bar in D = 5_μ_m. **g, h**. VGLUT1 count in adult *Tmem16f* cKO and control animals in superficial lamina I (**g**) and deep lamina III (**h**). LI VGLUT1 count mean difference 201.33 [95.00% CI 110.90, 292.00]; LIII VGLUT1 count mean difference 56.61 [95.00% CI −13.81, 118.42]. FOV = 31_μ_m x 94_μ_m. N-numbers as indicated. Black and red data points indicate females and males respectively.

As synaptic engulfment was also observed in the deeper laminae we re-analysed the data by cropping the original images to only contain LI or LIII to assess potential differences across laminae (Figure 2e). Surprisingly, VGLUT1 density in LIII was unaltered in contrast to LI (Figure 2g-h), suggesting that the changes observed are mainly driven by an increase in superficial VGLUT1 terminal projections (unpaired mean difference superficial/LI VGLUT1 count: 201.33 [95.00% CI 110.90, 292.00], deep/LIII VGLUT1 count: 56.61 [95.00% CI −13.81, 118.42]).

To investigate whether the superfluous VGLUT1 terminals indeed form synaptic contacts, we also co-stained spinal cord sections from *Tmem16f* cKO and control animals with HOMER, which marks postsynaptic densities. Indeed, we found colocalization of VGLUT1-positive Thy1-GFP labelled A-fibre terminals with HOMER were increased in *Tmem16f* cKO animals, suggesting that a surplus of functional VGLUT1 positive synaptic contacts are formed in the superficial dorsal horn of *Tmem16f* cKO animals (Figure 2—figure supplement 1).

Thus, targeted deletion of microglial *Tmem16f*, and consequently impaired phagocytosis specifically disrupted the normal developmental pruning of excitatory afferent A-fibre projections in the superficial dorsal horn.

### Dorsal horn sensory neurons in adult microglial Tmem16f cKO mice are less responsive to dynamic tactile stimulation of the skin, but have larger receptive fields

We reasoned that the reduced A-fibre pruning due to impaired neonatal microglial phagocytosis in *Tmem16f* cKO mice would alter behavioural sensitivity to hindpaw tactile stimulation. The hindlimb withdrawal reflex was measured in response to brushing of the plantar surface of the paw. *Tmem16f* cKO mice showed a greater number of withdrawals in response to dynamic brushing than controls (unpaired mean difference 0.83 [95% CI 0.072, 1.57]), but not to static vF stimulation (Figure 3, Figure 3—figure supplement 1). *Tmem16f* cKO animals therefore displayed cutaneous hypersensitivity specifically towards dynamic low-threshold tactile stimulation.

**Figure 3.**
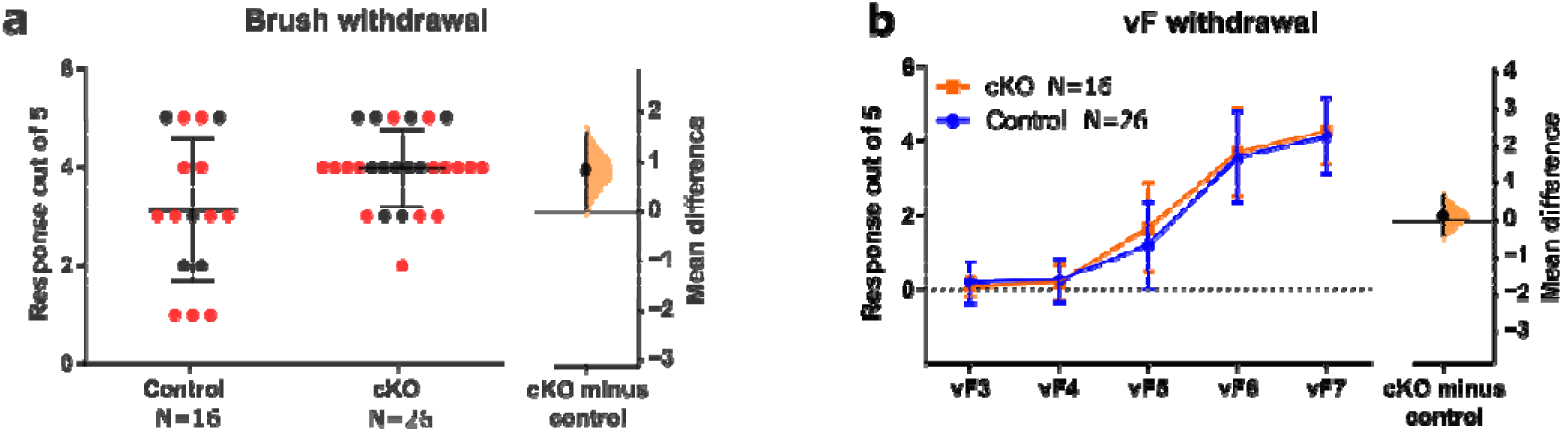
Neonatal *Tmem16f* deletion in microglia increases dynamic brush sensitivity. **a**. Brush withdrawal response for *Tmem16f* cKO animals are higher than control animals, mean difference 0.83 [95% CI 0.072, 1.57]. **b**. vF withdrawal response did not differ between *Tmem16f* cKO and control animals, mean difference 0.12 [95% CI −0.39, 0.65]. N-numbers as indicated. Black and red data points indicate females and males respectively.

We hypothesised that excessive A-fibre terminals in dorsal horn might lead to increased dorsal horn activity which could underlie the behavioural hypersensitivity. To test this, we used *in vivo* single unit extracellular recordings in the dorsal horn of anaesthetised *Tmem16f* cKO and control mice. We recorded from adapting and non-adapting wide dynamic range neurons (WDR) in the deep dorsal horn, which according to the criteria by Lee et al. 2019 ^32^ are presumed to be excitatory and inhibitory neurons respectively. Of 30 cells recorded from control mice, 8 were non-adapting and 22 were adapting. Of 28 cells recorded from *Tmem16f* cKO mice, 5 were non-adapting and 23 were adapting. This is consistent with the expected ratio of 1:2 of inhibitory to excitatory cells as reported in previous studies ^32–34^ (P=0.56 for control and P=0.11 for cKO in Binomial test against expected ratio). We quantified spinal neuronal response to 1) dynamic innocuous tactile stimuli (brush), and 2) static von Frey hair (vF) stimuli applied to the plantar hindpaw as well as mapped dynamic touch-sensitive receptive field areas on the plantar hindpaw.

We reasoned that excessive A-fibre presence in the dorsal horn would increase their spatial connectivity with postsynaptic neurons, leading to larger dorsal horn receptive field areas. Consistent with increased A-fibre synaptic contacts, mean brush receptive field sizes for adapting neurons were increased by almost 50% (unpaired mean difference 13.7 [95.00% CI 3.3, 27.2]), with a notable subpopulation of cKO neurons expanding their receptive fields to cover the entire, or majority of the plantar surface (Figure 4a, b). The expansion of receptive field area was accompanied by a reduced number of spikes in response to both brush and vF stimulation in *Tmem16f* cKO adapting neurons (unpaired mean difference brush: −11.4 [95% CI −19.2, −4.5], vF: −6.4 [95% CI −8.5, −4.5]) as well as a reduction in spontaneous activity (Figure 4c-f, Figure 4—figure supplement 1a).

**Figure 4.**
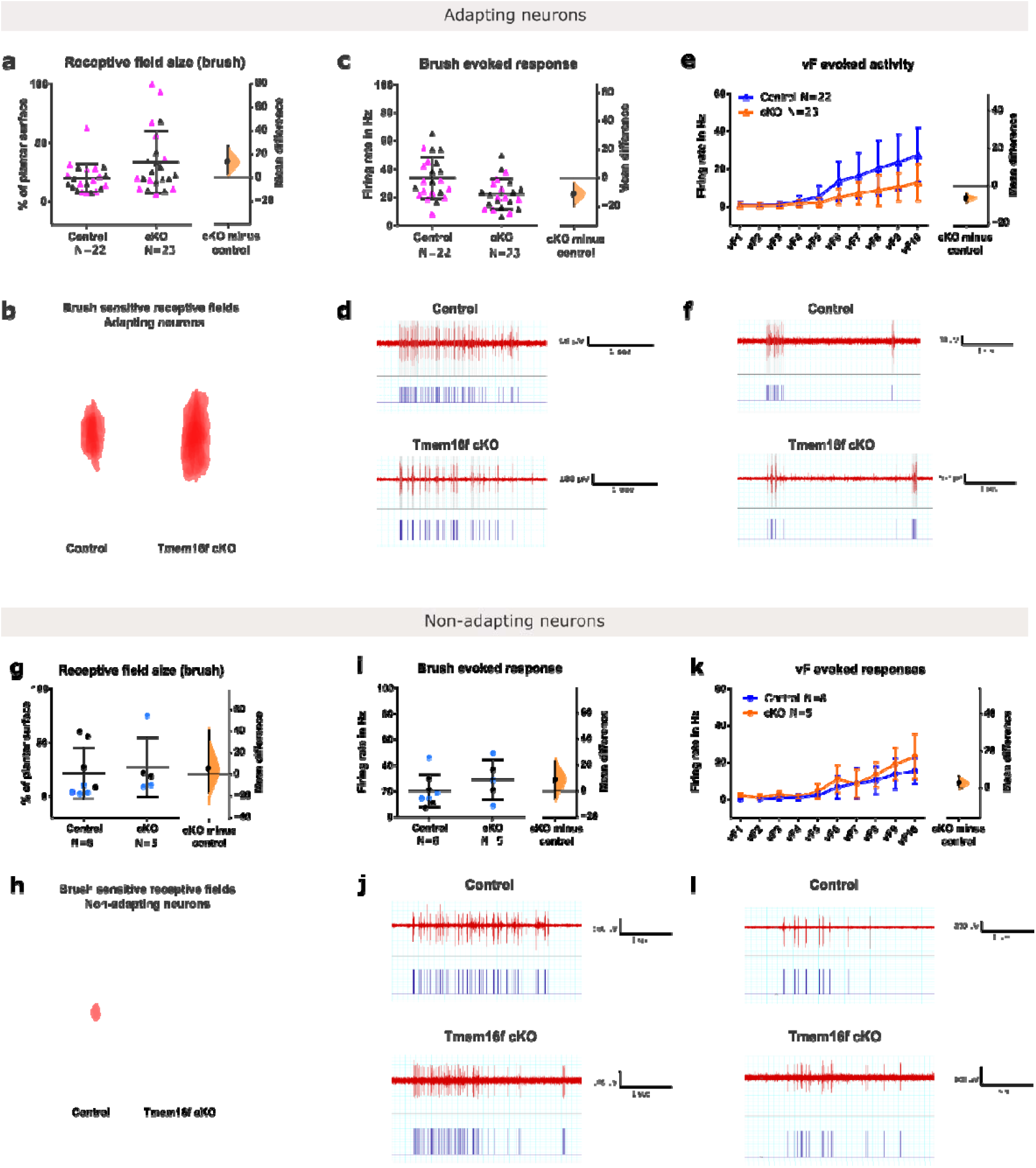
Neonatal *Tmem16f* deletion in microglia decreases evoked activity, but increases brush receptive field size. **a**. Brush receptive field sizes was increased for adapting neurons in *Tmem16f* cKO, mean difference 13.7 [95.00% CI 3.3, 27.2]. **b**. Overlay of receptive fields in **a**. **c**. Brush evoked response was decreased for adapting neurons in *Tmem16f* cKO, mean difference - 11.4 [95% **CI −19.2, −4.5]**. **d**. Example firing trace of adapting cell from control and cKO animals to brush stimulation with raster plots underneath. **e**. vF evoked activity was decreased for adapting neurons in *Tmem16f* cKO, mean difference −6.4 [95% CI −8.5, −4.5]. **f**. Example firing trace of an adapting cell from control and cKO animals to vF stimulation with raster plots underneath. **g**. Brush receptive field sizes was unchanged for non-adapting neurons in *Tmem16f* cKO, mean difference 5.53 [95.00% CI −15.91, 40.43]. **h**. Overlay of receptive fields in **g**. **i**. Brush evoked response was unchanged for non-adapting neurons in *Tmem16f* cKO, mean difference 8.62 [95.00% CI −5.78, 22.72]. **j**. Example firing trace of non-adapting cell from control and cKO animals to brush stimulation with raster plots underneath. **k**. vF evoked activity was decreased for non-adapting neurons in *Tmem16f* cKO, mean difference 2.74 [95.00% CI −0.074, 6.25]. **l**. Example firing trace of a non-adapting cell from control and cKO animals to vF stimulation with raster plots underneath. N-numbers as indicated. Black and colourful data points indicate females and males respectively.

In contrast, brush receptive field sizes and brush-evoked responses did not differ between control and *Tmem16f* cKO mice in non-adapting (inhibitory) neurons (Figure 4g-j), though an increase in vF-evoked responses in *Tmem16f* cKO animals was observed (mean difference 2.74 [95.00% CI −0.074, 6.25], F(1; 110) = 7.014, P = 0.0093) (Figure 4k, l). Spontaneous activity was also unaltered (Figure 4—figure supplement figure 1b).

## Discussion

In summary, we report that dorsal horn microglia have a distinct phenotype during the first postnatal week characterized by high phagocytic activity, which coincides with postnatal engulfment of A-fibre terminals in the dorsal horn, supporting a role of microglia in developmental dorsal horn remodelling. We further show that disruption of microglial function by targeted deletion of *Tmem16f* during early postnatal life impairs microglia mediated A-fibre refinement in the dorsal horn leading to long-term changes in dorsal horn function and behaviour that persist into adulthood. Together, our data suggests that microglia mediated refinement of A-fibres during the early postnatal period is critical to both normal dorsal horn development and appropriate spatial encoding of dynamic touch.

In our study we looked at a subset of VGLUT1 positive A-fibres, which are predominantly low-threshold, myelinated Aβ mechanoreceptors ^24,35^, and primarily transduce innocuous mechanical stimulation of the skin, such as stroking and brushing. In adults, the superficial laminae is occupied by both A- and C-fibre projections, but the during the neonatal period A-fibre inputs dominate the superficial dorsal horn, as nociceptive C-fibre inputs are weak and only start to strengthen between P5-P10 ^36,37^. Functionally, this results in lower cutaneous mechanical thresholds for activity in the superficial laminae, which together with immature local and descending inhibition leads to exaggerated reflex behaviour in neonates ^5–7,38^. This early sensitivity to low threshold stimuli might be particularly important during the neonatal period for maternal bonding, as skin to skin contact after birth was shown to reduce neonatal stress and pain response and improve mother-infant interaction ^39^. Therefore, appropriate A-fibre maturation might play an important role in overall infant development.

Microglia cells show the strongest increase in density over the first postnatal week which coincides with a decrease in phagocytic activity. Hammond et al. 2019 revealed a specialised phagocytic population of microglia at P4/5 in the brain, while our data suggests that the peak of microglial phagocytic activity might be later in the spinal cord around P10. Moreover, dorsal horn microglial density observed in our study appears to be higher than in various brain regions studied at comparable ages ^21,40^ (Table 1). Together, this suggests that dorsal horn microglia follow a different developmental trajectory than brain microglia.

**Table 1.** Microglial density across different CNS regions (in rats). Estimated numbers from the graphs in respective paper, with male and female microglia counts pooled.

| Paper | Brain area | Age | Density per $\mu\text{m}^3$ |
| --- | --- | --- | --- |
| Schwarz et al. 2012<sup>21</sup> | Parietal cortex | P4 | $2.23 \times 10^{-7}$ |
| | Amygdala | P4 | $4.77 \times 10^{-6}$ |
| | CA1 of hippocampus | P4 | $4.88 \times 10^{-7}$ |
| | Paraventricular nucleus | P4 | $1.37 \times 10^{-6}$ |
| Perez-Pouchoulen et al. 2015<sup>40</sup> | Cerebellum | P5 | $3 \times 10^{-6}$ |
| Present study (Figure 1—figure supplement 1d) | Superficial dorsal horn | P3 | $5 \times 10^{-6}$ |

While microglial CD68 volume increased postnatally, the amount of microglial phagocytic cup numbers decreased. Phagocytic cups are relatively large structures involved in phagocytosing cell bodies ^41^, and their reduction is consistent with the decline in apoptotic cell numbers postnatally. This shows that the postnatal increase in CD68 cannot be due to removal of apoptotic cell bodies but rather due to the refinement of neuronal connections.

Dorsal horn microglia phagocytose A-fibre projections both in the superficial (laminae I-II) and deeper (laminae III-IV) laminae during normal postnatal development. The data here shows that most of the A-fibre engulfment occurs before P10 and decreases thereafter. The overall amount of A-fibre engulfment is likely underestimated, as reporter mice crossed with the *Vglut1*-Cre line used here only express tdT in a subset of A-fibres ^24^. Further, the developmental upregulation of *Vglut1* expression ^26,27^ means that engulfment will be especially underestimated in younger animals, as their A-fibres might be present but not yet expressing tdT. In the brain, it has been shown that VGLUT1 protein expression at P0 is only 6.6% of the adult level, reaching 47% by P10 and 92% by P20 ^27^. Therefore, the fall in A-fibre engulfment over the first postnatal week is likely much sharper in reality than Figure 1 g, h suggests.

Although microglial CD68 volume peaks at P10, the engulfed VGLUT1-tdT volume seems to peak earlier at P7. While the percentage of lysosome volume occupied by VGLUT1-tdT is about 15% in the superficial laminae and 40% in the deep laminae at P7, by P10 it has become less than a quarter of that. It is possible that a slowing rate in engulfment of VGLUT1-tdT labelled fibres between P10 and P7 allows VGLUT1-tdT to be degraded in the lysosomes at a faster rate than new VGLUT1-tdT material is engulfed, thus causing a reduction of VGLUT1-tdT at P7 before the reduction in lysosome volume follows suit by P10. It is also possible that developmental events at P10 require additional microglial phagocytosis of materials other than A-fibres.

Neonatal *Tmem16f* cKO increased the number of Thy1-GFP positive VGLUT1 synapses in the superficial dorsal horn in adults, suggesting that *Tmem16f* function is necessary for microglia mediated refinement of A-fibres. However, increase in presynaptic terminals seems to predominantly affect lamina I, but not lamina III, even though developmental engulfment of A-fibres was observed both in LI-II and LIII-IV, suggesting that either only the engulfment of superficial synapses is dependent on *Tmem16f* or alternatively that superfluous synapses in the deeper laminae were more efficiently compensated and removed at later stages before animals reached adulthood.

The density of VGAT presynaptic terminals was also unaltered in *Tmem16f* cKO animals. This suggests that *Tmem16f* mediated engulfment is synapse specific. A recent study in the hippocampus suggested that complement mediated synapse elimination by microglia was targeted to VGLUT2, but not VGLUT1 or VGAT synapses instead ^42^. Together with our data this suggest that developmental synapse elimination is both specific to synapse identity and location.

Consistent with the increase in dorsal horn VGLUT1, *Tmem16f* cKO increased brush receptive field size in adapting/excitatory neurons, which mirrors the large receptive field sizes in immature neonatal animals ^5^. During the neonatal period A-fibre inputs dominate the superficial dorsal horn ^36,37^ and hindpaw receptive fields are initially large, reducing in size over the first two postnatal weeks ^5^. Functionally, this results in lower cutaneous mechanical thresholds, which together with immature local and descending inhibition leads to exaggerated reflex behaviour in neonates ^5–7,38^. Both enlarged receptive field sizes and hypersensitivity were observed in Tmem16f cKO animals. This is unlikely due to local disinhibition ^43,44^, as spontaneous and evoked activity of adapting WDR neurons were not increased, suggesting that the larger receptive field sizes reflect a lack of primary afferent refinement instead.

Consistent with the increased A-fibre input and brush receptive field size, which likely affects spatial summation of dynamic inputs, *Tmem16f* cKO increased behavioural sensitivity to brush. However, counterintuitively, the increase in A-fibre projections and behavioural sensitivity was accompanied by a reduction in evoked activity for brush and vF as well as spontaneous activity in adapting neurons.

How does decreased evoked activity result in higher reflex sensitivity? One possibility is that the activation pattern of WDR neurons may be more important than firing rate per se for the coding of sensory information. Although the activity of individual neurons is decreased, potentially more neurons are activated in total due to the lack of A-fiber refinement. Therefore, the activation of WDR neurons on a population level might be a better predictor of the spinal reflex response. A potential mechanism underlying this effect is homeostatic synaptic downscaling, which maintains homeostasis by reducing neuronal firing rate when the overall network activity is increased ^45^. Our data shows that activity recorded from single neurons do not always positively correlate with behavioural changes and need to be interpreted with care.

In contrast to the adapting neurons, little change was observed for non-adapting/inhibitory neurons, apart from an increase in vF-evoked activity. This is likely due to the low sample size, as fewer non-adapting/inhibitory neurons were recorded. The increased firing rate of non-adapting/inhibitory neurons during vF-evoked activity would likely decrease the activity of adapting/excitatory neurons and thus contribute to the decreased overall activity observed in cKO animals.

We propose therefore that disrupting microglial function with *Tmem16f* cKO causes an increase in VGLUT1 positive A-fibre terminals in adult animals, potentially due to failed pruning of A-fibres in the neonatal period. This excess of peripheral input might preferentially target excitatory neurons which leads to larger receptive field sizes thereof, and potentially activates larger numbers of excitatory neurons upon peripheral stimulation. In compensation, through homeostatic scaling or increased A-fibre input onto non-adapting/inhibitory neurons, inhibitory activity onto excitatory neurons is increased, decreasing the firing rate of individual neurons, but not necessarily the total output as a population. This is also consistent with increased behavioural sensitivity in *Tmem16f* cKO animals.

### Limitations

Several caveats need to be considered when interpreting the data. In our experiments we used a CNS-wide KO of microglial *Tmem16f*. Therefore, we cannot rule out cortical contributions to the observed changes in the spinal cord – for example, top-down facilitation of inhibitory inputs or inhibition of excitatory inputs could also explain the decreased firing rate in adapting and non-adapting neurons in *Tmem16f* cKO animals. However, a purely top-down mediated overall inhibition of dorsal horn activity would not explain the increase in receptive field size following *Tmem6f* 1 cKO, though it could additionally contribute to the changes observed. Therefore, at least part of the observed changes must be mediated by changes in peripheral afferent input to the dorsal horn.

Furthermore, it is possible that *Tmem16f* cKO will also affect the engulfment and removal of apoptotic cells by microglia, which contribute to the behavioural and functional changes seen in adulthood. However, the mechanisms involved in engulfing whole cell bodies vs synaptic material are likely distinct (in our study CD68 volume was not correlated with apoptotic clearance), and therefore *Tmem16f* cKO might affect microglial pruning without affecting the phagocytosis of apoptotic cells.

We used VGLUT1 as a marker for presynaptic boutons of A-fibres as well as the driver of tdT expression. However, it has been shown that cortico-spinal projections also contribute to VGLUT1 synapses throughout the dorsal horn ^33^. Therefore, we cannot exclude that the behavioural and functional effects seen in the *Tmem16f* cKO animals are due to disruption of cortico-spinal refinement. However, during early postnatal development, corticospinal projections do not enter in the grey matter lumbar segment until P7, while peripheral A-fibre afferents reach the lumbar dorsal horn by embryonic day (E)14 ^46,47^. Therefore, we can conclude that at least prior to P7, where the majority of VGLUT1-tdT engulfment was observed, the engulfment of VGLUT1-tdT labelled fibres are indeed afferent A-fibres.

An important open question is what determines the engulfment or survival of synapses and fibres in the dorsal horn. It is known that microglia can sense and respond to neuronal activity ^48–51^. For example, microglia increase process motility both to neuronal hyper- and hypoactivity ^49^, and in the brain, microglial engulfment seems to preferentially target weak synapses ^12,13^. Given that perturbation of neuronal activity can indeed impair A-fibre refinement ^8,9^, neuronal activity seems a likely signal for microglial engulfment in the dorsal horn. Several molecules expressed by neurons and microglia have been identified as “eat me” and “keep me” signals ^12,13,52,53^, among which members of the complement cascade, e.g. C1q, have been shown to tag synapses for microglial engulfment both in the brain and the spinal cord ^12,17^. Whether C1q also plays a role in dorsal horn remodeling remains to be investigated.

## Conclusions

In summary, we have shown that dorsal horn microglia phagocytose A-fibres during normal postnatal development and that disruption of microglial function can lead to long-term structural and functional changes in the dorsal horn and behavioural changes towards dynamic touch. This has important implications for perinatal care of infants.

Low-threshold stimuli such as skin-to-skin contact have been shown to reduce infant stress and improve infant-mother interaction and maternal bonding ^39^. Therefore, appropriate maturation of A-fibres might play an important role in overall infant development. In addition, previous research have shown that peripheral injury during the first postnatal week (but not after P10) cause hyperalgesia upon re-injury in adulthood, and that this could be prevented with intrathecal minocycline injection, which is a non-specific microglial inhibitor ^54,55^. It is tempting to speculate that injury alters microglial pruning of A-fibres as described here which could underlie part of the long-term changes seen following injury. Thus, the role of microglia in normal and aberrant neonatal development could be investigated for therapeutic potential in perinatal care.

## Materials and Methods

| Key Resources Table |  |  |  |  |
| --- | --- | --- | --- | --- |
| Reagent type (species) or resource | Designation | Source or reference | Identifiers | Additional information |
| Genetic reagent (*M. musculus*) | *Ai9*; *R26*<sup>LSL-Ai9/+</sup> | Jackson Laboratory | Stock # 007909<br>RRID:IMSR_JAX:007909 | PMID: 20023653 |
| Genetic reagent (*M. musculus*) | *Ai9*; *R26*<sup>LSL-Ai9/+</sup> | Jackson Laboratory stock | Stock # 007905<br>RRID:IMSR_JAX:007905 | PMID: 20023653 |
| Genetic reagent (*M. musculus*) | *Slc17a7*-IRES2-Cre; *Vglut1*<sup>Cre/+</sup> | Jackson Laboratory stock | Stock # 023527<br>RRID:IMSR_JAX:023527 | PMID: 25071457 |
| Genetic reagent (*M. musculus*) | *Cx3cr1*-CreERT2-YFP; *Cx3cr1*<sup>CreER/+</sup> | Jackson Laboratory stock | Stock # 021160<br>RRID:IMSR_JAX:021160 | PMID: 35045285 |
| Genetic reagent (*M. musculus*) | *Thy1*-EGFP-M; *Thy1*<sup>eGFP</sup> | Jackson Laboratory stock | Stock # 007788<br>RRID:IMSR_JAX:007788 | PMID: 11086982 |
| Genetic reagent (*M. musculus*) | *Tmem16f*-floxed; *Tmem16f*<sup>fl/fl</sup> | PMID: 27332874 |  | Dr Paul Heppenstall (SISSA, Trieste, Italy) |
| Antibody | anti-Caspase-3 (Rabbit monoclonal) | Cell Signalling | Cat# Asp175, 5A1E | IF (1:100) |
| Antibody | anti-CD68 (Mouse monoclonal) | BIO-RAD | Cat# MCA341R | IF(1:500) |
| Antibody | anti-CD68 (Rabbit polyclonal) | Abcam | Cat# ab125212 | IF(1:500) |
| Antibody | anti-GFAP<br>(Mouse,<br>monoclonal) | Sigma | Cat# G3893 | IF(1:2000) |
| Antibody | anti-Iba1<br>(Rabbit,<br>polyclonal) | Wako | Cat# 019-19741 | IF(1:1000) |
| Antibody | anti-Iba1<br>(Goat, polyclonal) | Abcam | Cat# ab5076 | IF(1:1000) |
| Antibody | anti-NeuN<br>(Mouse,<br>monoclonal) | Millipore | Cat# MAB377 | IF(1:2000) |
| Antibody | anti-VGAT<br>(Mouse,<br>monoclonal) | Synaptic<br>Systems | Cat# 131011 | IF(1:2000) |
| Antibody | anti-VGAT<br>(Rabbit,<br>polyclonal) | Synaptic<br>Systems | Cat# 131002 | IF(1:2000) |
| Antibody | anti-VGLUT2<br>(Guinea pig,<br>polyclonal) | Millipore | Cat# AB2251-I | IF(1:2000) |
| Antibody | anti-Synaptophysin<br>(Mouse,<br>monoclonal) | Abcam | Cat# Ab8049 | IF(1:2000) |
| Antibody | anti-Synaptophysin<br>(Guinea pig,<br>polyclonal) | Synaptic<br>Systems | Cat# 101004 | IF(1:2000) |
| Antibody | anti-goat Cy2<br>(Donkey<br>polyclonal) | Jackson<br>ImmunoResea<br>rch | Cat# 705-225-147 | IF(1:500) |
| Antibody | anti-goat Cy3<br>(Donkey<br>polyclonal) | Jackson<br>ImmunoResea<br>rch | Cat# 705-165-147 | IF(1:500) |
| Antibody | anti-goat Cy5<br>(Donkey<br>polyclonal) | Jackson<br>ImmunoResea<br>rch | Cat# 705-175-147 1 | IF(1:500) |
| Antibody | anti-goat Alexa 488 (Donkey polyclonal) | Jackson ImmunoResearch | Cat# 705-545-003 | IF(1:500) |
| Antibody | anti-guinea pig Cy3 (Donkey polyclonal) | Jackson ImmunoResearch | Cat# 706-165-148 | IF(1:500) |
| Antibody | anti-guinea pig Alexa 647 (Donkey polyclonal) | Millipore | Cat# AP193SA6 | IF(1:500) |
| Antibody | anti-mouse Cy2 Jackson (Donkey polyclonal) | Jackson ImmunoResearch | Cat# 715-225-150 | IF(1:500) |
| Antibody | anti-mouse Cy3 Jackson (Donkey polyclonal) | Jackson ImmunoResearch | Cat# 715-165-150 | IF(1:500) |
| Antibody | anti-mouse Cy5 (Donkey polyclonal) | Jackson ImmunoResearch | Cat# 715-175-150 | IF(1:500) |
| Antibody | anti-rabbit Cy2 (Donkey polyclonal) | Jackson ImmunoResearch | Cat# 711-225-152 | IF(1:500) |
| Antibody | anti-rabbit Alexa 594 (Donkey polyclonal) | Jackson ImmunoResearch | Cat# 711-585-152 | IF(1:500) |
| Antibody | anti-rabbit Alexa 647 (Donkey polyclonal) | Millipore | Cat# AP182SA6 | IF(1:500) |
| Antibody | anti-rabbit-biotinylated (Horse polyclonal) | Vector | Cat# BA11-00 | IF(1:500) |
| peptide, recombinant protein | Streptavidin-594 | Life Technologies | Cat# S32356 | IF(1:500) |
| Software, algorithm | ImageJ; FIJI | PMID:22930834, PMID:29187165 | RRID:SCR_003070 |  |
| Software, algorithm | Volocity | PerkinElmer | RRID:SCR_002668 |  |
| Software, algorithm | LabChart 7 | ADInstruments |  |  |
| Software, algorithm | Graphpad prism | GraphPad Software, Inc. | RRID:SCR_002798 |  |
| Software, algorithm | Estmationstats.com (web application) | PMID: 31217592 | RRID:SCR_018321 | Free web application |
| other | Carbostar-1 (micro-electrode) | Kation Scientific |  |  |

### Animals

Sprague Dawley rats of both sexes were used for experiments in Figure 1—figure supplement 1d-h and Figure 1—figure supplement 2. Transgenic mice on C57BL/6J background of both sexes were used in all other experiments.

Experiments used the following transgenic mouse lines:

1. Ai9 / Rosa26-CAG::loxP-STOP-loxP-tdTomato-WPRE (Jackson Laboratory stock 007909 & 007905)
2. *Slc17a7*-IRES2-Cre (Jackson Laboratory stock 023527)
3. *Cx3cr1*-CreERT2-YFP (Jackson Laboratory stock 021160)
4. *Thy1*-EGFP-M (Jackson Laboratory stock 007788)
5. *Tmem16f*-floxed (flx) animals (Batti et al., 2016)

For visualisation of A-fibres, *Slc17a7*-IRES2-Cre (*Vglut1*-Cre) males (JAX stock no. 023527) were crossed with Ai9 females (JAX stock no. 007909) to obtain animals that expressed the tdTomato fluorophore under the *Vglut1* promoter (*Vglut1*^*Cre/+*^; *R26*^*LSL-Ai9/+*^).

To generate tamoxifen inducible microglia-specific *Tmem16f* knock-out mice (*Tmem16f* cKO), Cx3cr1-CreER-YFP (JAX stock no. 021160) mice were crossed to *Tmem16f*-flx animals (generated by P. Heppenstall, see Batti et al. 2016), as well as Ai9 (JAX stock no. 007905) and *Thy1*-EGFP-M (JAX stock no. 007788).

Experimental animals were heterozygous for *Cx3cr1*-CreER-YFP (^CreER/+^), homozygous for mutant conditional allele *Tmem16f*-flx (^fl/fl^), and carrying Ai9 (*R26*^*LSL- Ai9*^) and *Thy1*-eGFP (^eGFP^) alleles (zygocity was not determined for *R26*^*LSL-Ai9*^ and *Thy1*^*eGFP*^). This produced the following genotype: *Cx3cr1*^*CreER/+*^; *Tmem16f*^*fl/fl*^; *R26*^*LSL-Ai9*^; *Thy1*^*eGFP*^. Control animals were homozygous for the wild type *Tmem16f* allele: *Cx3cr1*^*CreER/+*^; *Tmem16f*^*+/+*^; *R26*^*LSL-Ai9*^; *Thy1*^*eGFP*^.

Both females and males were used. No sex differences were expected and animals of both sexes were pooled together for analysis, but data points are presented as black (female) or red/magenta/blue (male) to indicate the sexes. Numbers of animals used for each experiment are indicated in the figures. For a table with detailed species, ages, sexes, and numbers of animals used in each experiment, please see supplementary Table 1. All procedures were carried out in accordance with the guidelines of the UK Animals (Scientific Procedures) Act 1986 and subsequent amendments.

### Drugs

4-hydroxytamoxifen (4-HT) was dissolved at 1mg/ml in corn oil, and 50μl was injected intragastrically per pup daily on three consecutive days from P1-3, following a previously described protocol ^56^. Both control and experimental animals (*Tmem16*^*+/+*^ and *Tmem16f*^*fl/fl*^ respectively, see animals section above) received 4-HT injections to control for any effects of 4-HT itself. The dam was given a protein enriched diet a few days before and following delivery to aid milk production and pup survival.

### Immunohistochemistry

Animals were overdosed with pentobarbital and transcardially perfused with saline followed by ice-cold 10% formalin. The sciatic nerve was exposed and traced to locate L4 & L5 dorsal root ganglia (DRG) and the corresponding region of the lumbar spinal cord was dissected and post-fixed in 10% formalin overnight, followed by immersion in 30% sucrose until they sank. 50μm free-floating spinal cord sections were cut on the microtome with every 2nd section collected.

Tissue sections were washed 3 × 10 min in PBS and then incubated in blocking solution (10% donkey serum, 0.2% Triton X-100 in PBS) for 2.5h at room temperature. The sections were then incubated with primary antibodies at 4°C overnight followed by secondary antibodies at room temperature for 2h, both diluted in 3% blocking solution (3% donkey serum, 0.2% Triton X-100 in PBS) (for list of antibodies and their respective concentrations used, see Key Resources Table). Samples were mounted in Fluoromount Aqueous Mounting Medium (Sigma) or ProLong™ Diamond Antifade Mountant (Thermo Fischer), if the tissue contained endogenous fluorophores.

### Image acquisition and analysis

Confocal z-stacks were taken with a Zeiss LSM880 confocal microscope or Yokogawa CSU22 spinning disk microscope using a 20× water immersion objective (NA 1.0) for imaging of A-fibres and 63× oil immersion objective (NA 1.4) followed by analysis in Fiji software. Details on microscope settings can be found in supplementary >Table 1 and in the metadata of example images online at https://github.com/Yajing826/A-fibre-engulfment. Only intact sections with an even stain were analysed, and at least 6 sections were imaged and analysed per animal to reduce variability for all figures (apart from S3, where 1-3 sections were analysed per animal).

Cell counts and phagocytic cup counts were performed manually on confocal images in Fiji or Volocity, with the experimenter blinded to the age groups. Phagocytic cups were defined as any rounded structure at the end of a microglial process.

A-fibre engulfment by microglia and synapse density were analysed with automated batch processing in Fiji using the 3D-ROI manager plugin and custom written macros ^57–60^. For A-fibre engulfment, each of the channels containing staining for microglia, lysosomes, or A-fibres were binarized and the volume of their overlap measured. For synapse density measures, the channel containing synaptic stain was binarized and segmented, following which volume and object numbers were recorded. Macro scripts for the automated analysis are available online at https://github.com/Yajing826/A-fibre-engulfment.

### Behaviour

Behavioural testing was carried out on adult mice of both sexes between 3-4 months old, with the experimenter blinded to animal genotype/treatment. Animals were placed on a mesh platform (Ugo Basile) within individual transparent plastic chambers (6cm x 6cm x 12cm) for sensory testing of the plantar surface of the hind paw. Habituation and testing happened over five consecutive days. Animals were habituated to the testing environment for 1h per day on the first two days within individual plexicon chambers on the mesh platform. On the remaining days, animals were habituated for 30 min before being tested on each day. Brush response and von-Frey (vF) threshold were determined on the 3rd day, while repeated vF response testing was spread over the remaining 2 days to avoid sensitisation. Number of withdrawal reflexes were scored in each case, where only a rapid paw lifting was scored as a reflex. Animals were allowed to rest at least 20 seconds between each stimulus. For brush response, a fine brush (Pro Arte, series 007, size 2) was moved over the plantar surface of the hind paw from heel to toe over a 2 second period ^61^. This was repeated five times and the number of withdrawal reflexes out of five was recorded. vF threshold was assessed using the simplified up and down method ^62^. Filaments were aimed at the region between the foot pads. Force was applied until the filament bent, and held in place for 2 seconds.

To generate a response curve to vF stimulation, repeated vF response was recorded by applying each of filaments no. 3 - 7 (0.04g - 0.6g) five times on the plantar surface, directed at the region between the foot pads. The sequence of vF filaments was randomised. Number of withdrawal reflexes out of five times was recorded.

### In vivo extracellular recording

Animals subjected to behavioural testing were reused in electrophysiological recordings. Experimenter was blinded to animal genotype/treatment. All recordings were performed on adult mice (3-4 months old) of both sexes in the deep dorsal horn. Cells were not recorded beyond 550 μm depth from the surface of the spinal cord (see Figure 4—figure supplement 1c for depth of all recorded neurons). 3-5 mice were used per sex and treatment group. All experiments were carried out by the same experimenter to ensure consistency.

#### Animal preparation

Mice were anaesthetised with intraperitoneal urethane injection (10% in saline, 1.5g/kg). 100μl of 0.6 g/ml atropine and 200μl saline were injected subcutaneously to respectively counteract the mucus-driving side effect of urethane and to prevent dehydration. The animal was constantly monitored for depth of anaesthesia throughout the experiment and supplemented with 50μl (5μg) urethane as needed. 200μl of saline was supplemented every 2 hours. Body temperature of the animal was kept close to 37°C with a heating pad throughout.

After cessation of reflexes, a tracheotomy was performed and a short plastic tube of about 1cm inserted to aid free breathing of the animal. The animal was then transferred onto a stereotactic frame and fixed with ear and hip bars. A laminectomy was carried out at vertebral level T13-L1 which corresponds to the spinal segments L4-L5 underneath. The spinal column was clamped for stability, the dura was removed, and the exposed spinal cord was covered with mineral oil to prevent drying.

#### Single unit extracellular recordings

A carbon micro-electrode (Carbostar-1, Kation Scientific) was lowered with a motorised manipulator (Scientifica) into the exposed spinal cord at a constant speed. A reference electrode was inserted into the back muscle close to the laminectomy for differential recording. Recorded neural activity was amplified 2000 times, and filtered for signals between 1kHz-10kHz (NL104 amplifier and NL125/6 band-pass filter modules from NeuroLog Digitimer). The signal was sampled at 20kHz and digitised using Powerlab 4/30 (ADInstruments). The trace was recorded and analysed in the software LabChart 7 (ADInstruments).

To isolate single neurons, the plantar surface of the animal’s hind paw was gently continuously stroked as a searching signal, while the electrode is being lowered through the dorsal horn of the spinal cord. Once a cell has been identified by equal amplitude of the spikes recorded, the brush receptive field of the cell was mapped out by a fine brush (Pro Arte, series 202, size 1, brush tip cut short to 7mm length x 1mm width).

Spontaneous activity was recorded for 10 min before and 5 min after stimulation. To record brush and von-Frey (vF) fibre evoked activity from single neurons, each stimulus was manually applied for 2 seconds over the receptive field of the cell and repeated 3 times, with a minimum of 10 second interval in between. (vF filament strength were as follows: 1 = 0.008g, 2 = 0.02g, 3 = 0.04, 4 = 0.07g, 5 = 0.16g, 6 = 0.40g, 7 = 0.60g, 8 = 1g, 9 = 1.7, 10 = 2g).

Cells were not recorded if they exhibited very high spontaneous firing rates that did not allow evoked activity to be clearly distinguished from spontaneous activity. Only wide dynamic range neurons responding both to brush and pinch stimulation were recorded. Animals were euthanised at the end of the experiment and the spinal cord was collected in neutral buffered 10% formalin (overnight) followed by 30% sucrose solution for re-use in immunohistochemistry.

#### Analysis & cell type categorisation

Analysis was carried out in the LabChart 7 software (ADInstruments). For spontaneous activity, firing rate was analysed over a 10min window prior to applying any stimuli. Firing rate for evoked responses were analysed over the first second of the stimulus duration and averaged over three trials.

Cells were divided into adapting and non-adapting groups based on their firing properties towards a threshold vF-stimulus ^32^. This threshold vF was defined as the first vF filament that evokes a firing rate of 10Hz or more. The response within the first second of vF application was analysed to calculate an adaptive ratio R, which was defined as

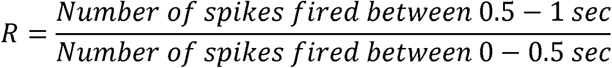

If a cell adapts rapidly to stimulation, one would expect R to be close to zero, as barely any spikes should be fired between 0.5 - 1 sec, however, if a cell is non-adapting and firing continuously, one would expect R to be close to 1. To decide the boundary between adapting and non-adapting cells we used k-means cluster analysis, which sorted the values into two groups that is equivalent to a boundary at R = 0.33. For the k-means clustering we included cells from a previous experiment.

### Statistical Analysis

Estimation statistics for the 95% confidence intervals (95% CI) of the mean difference were calculated on estimationstats.com ^63^ using 5000 samples of bias-corrected and accelerated bootstrapping. As bootstrapping is less accurate for small samples sizes ^64^, confidence intervals were only calculated for samples with N≥5. For Figure 1—figure supplement 1d, h and Figure 1—figure supplement 2 where N=4, one-way ANOVA was used with post-hoc comparisons carried out using Dunnett’s method.

Additionally, conventional null-hypothesis significance testing was carried out on estimationstats.com and GraphPad Prism 6 for all comparisons (significance level was set at α=0.05), which are listed in the Appendix 1—table 1.

Data are presented as mean ± SD in all figures. Where applicable, the effect size is presented as 95% CI of the mean difference on a separate but aligned axis. The mean difference is plotted as a dot on the background of its probability distribution, and the 95% confidence interval is indicated by the ends of the error bar. All values in text and figures are given with two decimals or rounded to two significant figures. N-numbers are as indicated in figures. For a comprehensive list with exact statistical values and analyses, see Appendix 1—table 1.

## Data availability

All data generated or analysed during this study are included in the manuscript and supporting files. Source files for all data are available at: https://www.ebi.ac.uk/biostudies/ under the accession number S-BSST609. Macro scripts for automated analysis in Fiji are available online at https://github.com/Yajing826/A-fibre-engulfment.

## Author Contributions

YX and SB performed the experiments, YX, SK, MF and SB designed the study, analysed and interpreted data and co-wrote the manuscript. AC, QH and RRJ developed the mouse model and prepared tissue for analysis. MS and PH developed the mouse model.

## Supplementary figures

**Figure 1—figure supplement 1.**
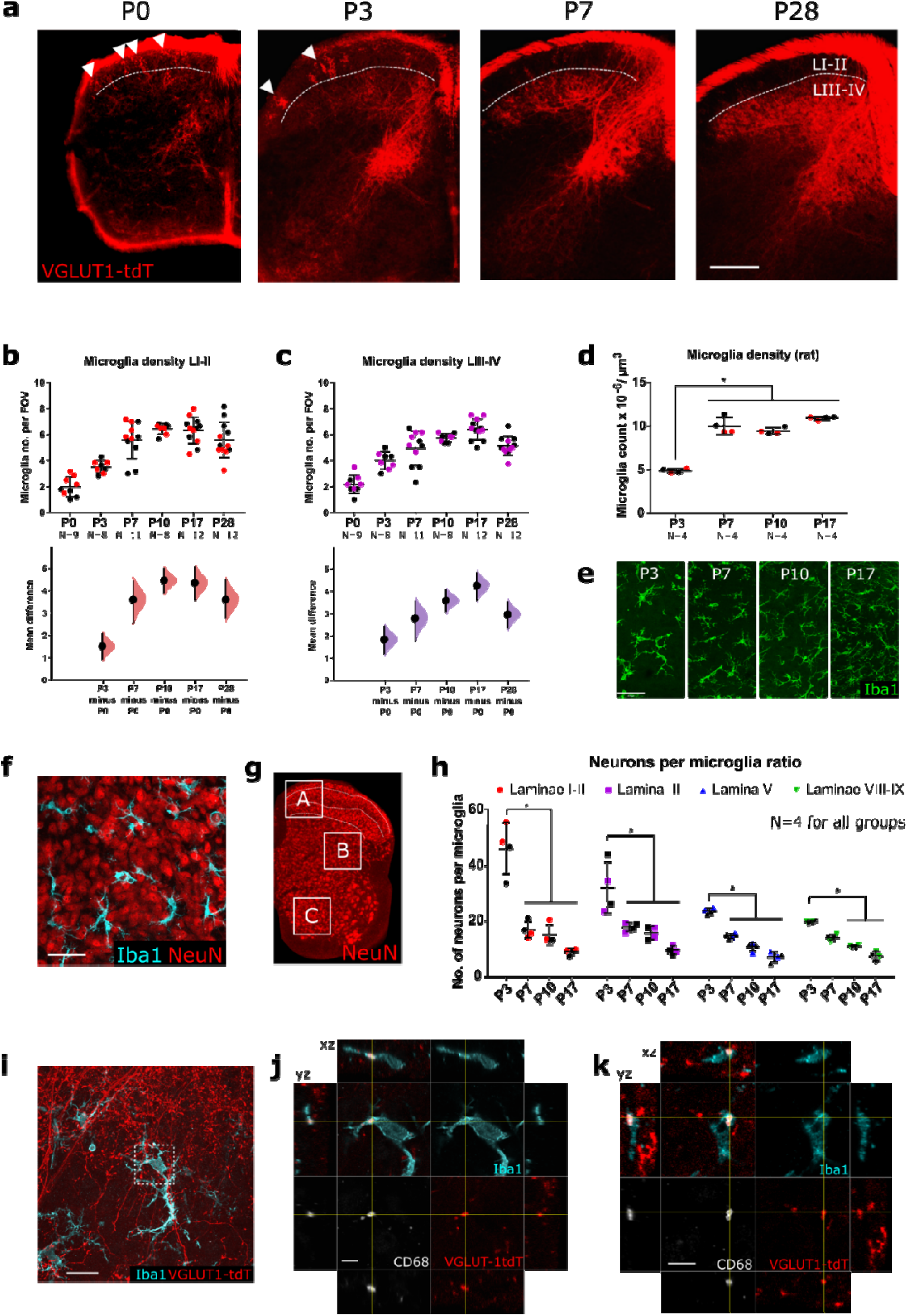
Microglial density changes over postnatal development. **a**. Representative images showing an increase in tdT-expression in the dorsal horn with age. Dashed white line indicates border between lamina II and lamina III, white arrow heads indicate tdT-fibres in the superficial laminae I-II. Scale bar = 200_μ_m. **b**. Microglial density increases over the postnatal period in both LI-II. P3 vs. P0: mean difference 1.53 [95% CI 0.97, 2.08]; P7 vs. P0: mean difference 3.61 [95% CI 2.58, 4.42]; P10 vs. P0: mean difference 4.46 [95% CI 3.91, 4.98]; P17 vs. P0: mean difference 4.35 [95% CI 3.60, 5.06]; P28 vs. P0: mean difference 3.61 [95% CI 2.79, 4.49]. Field of view (FOV) = 245_μ_m x 65_μ_m. **c**. Microglial density increases over the postnatal period in LIII-IV. P3 vs. P0: mean difference 1.83 [95% CI 1.21, 2.42]; P7 vs. P0: mean difference 2.79 [95% CI 1.80, 3.54]; P10 vs. P0: mean difference 3.58 [95% CI 3.09, 4.06]; P17 vs. P0: mean difference 4.24 [95% CI 3.60, 4.81]; P28 vs. P0: mean difference 2.96 [95% CI 2.35, 3.51]. Field of view (FOV) = 245_μ_m x 65_μ_m. **d**. Microglia density in rats increases significantly between P3 and P7, with P < 0:0001 for all ages compared to P3. **e**. Representative images of microglia cells in the dorsal horn (LI-III) stained for Iba1 (green) at postnatal days (P)3, P7, P10, P17. Scale bar = 50 _μ_m. **f**. Maximum projected confocal images from the dorsal horn (LI-III) at P3 with representative staining for microglia (Iba1, cyan) and neurons (NeuN, red). Scale bar = 50_μ_m. **g**. P7 spinal cord hemisection stained with neuronal marker NeuN (red), white boxes A, B, C indicate the approximate area of laminae I-III, lamina V, and laminae VIII-IX used for analysis. **h**. Ratio of neurons per microglia significantly decreases with age in laminae I-II, lamina III, lamina V, and laminae VIII-IX. **i**. Representative confocal image of microglial A-fibre engulfment in the spinal dorsal horn of P3 VGLUT1-tdt mice, stained for microglia (Iba1, cyan), microglial lysosomes (CD68, white) and endogenously fluorescent A-fibres (tdTomato, red). Scale bar = 20_μ_m. White inset box show location of higher magnification panels in **i**. **j**. High magnification image of microglial A-fibre engulfment in **h** stained for microglia (Iba1, cyan), microglial lysosomes (CD68, white) and endogenously fluorescent A-fibres (tdTomato, red). Cross-hairs show position of the xz and yz side-view panels. Scale bar = 5_μ_m. **k**. Further example of microglial A-fibre engulfment, colours and scale as indicated in **i**. N-numbers as indicated. Black and colourful data points indicate females and males respectively.

**Figure 1—figure supplement 2.**
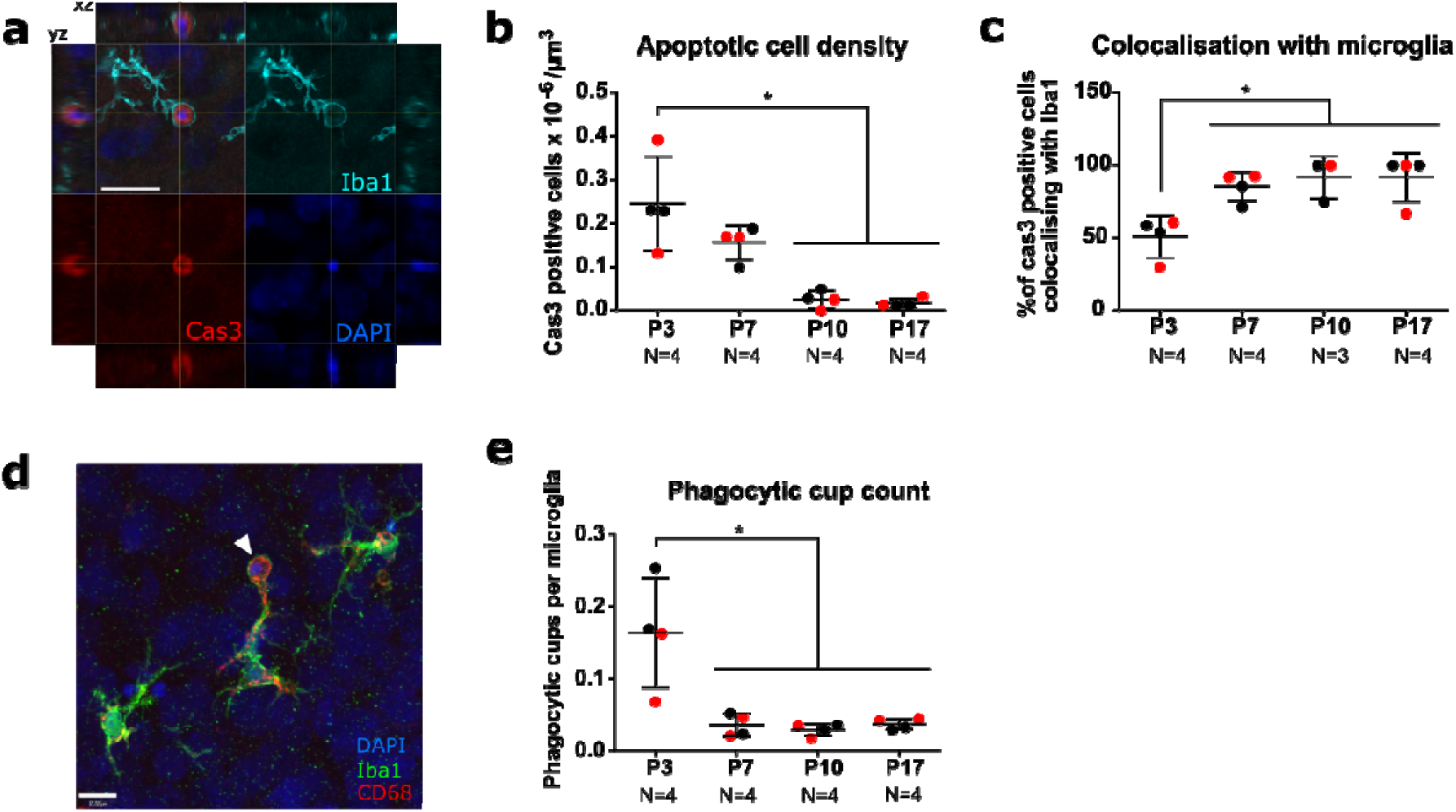
Apoptotic cell clearance over the postnatal period. **a**. Representative confocal image from the P3 dorsal horn of a microglia cell engulfing an apoptotic cell with a phagocytic cup. Cross-hairs show position of the xz and yz side-view panels. Microglial cell body is outside of the visible xy-plane. Image shows representative staining for Iba1 (cyan), Cas3 (red), DAPI (blue) at P3. Scale bar = 20_μ_m. **b**. Apoptotic cell numbers decrease significantly by P10, with P < 0.001 for P10 and P17 compared to P3. **c**. The percentage of Cas3 positive cells colocalising with microglia significantly increases with age, with P < 0.05 for all ages compared to P3. **d**. Representative image of microglia cell in the dorsal horn with phagocytic cup indicated by white arrowhead. Image shows representative staining for Iba1 (green), CD68 (red), DAPI (blue) at P3. Scale bar = 10_μ_m. **e**. Phagocytic cup count per microglia decreases significantly between P3 and P7, (P < 0.01 for all ages compared to P3). N-numbers as indicated. Black and red data points indicate females and males respectively.

**Figure 2—figure supplement 1.**
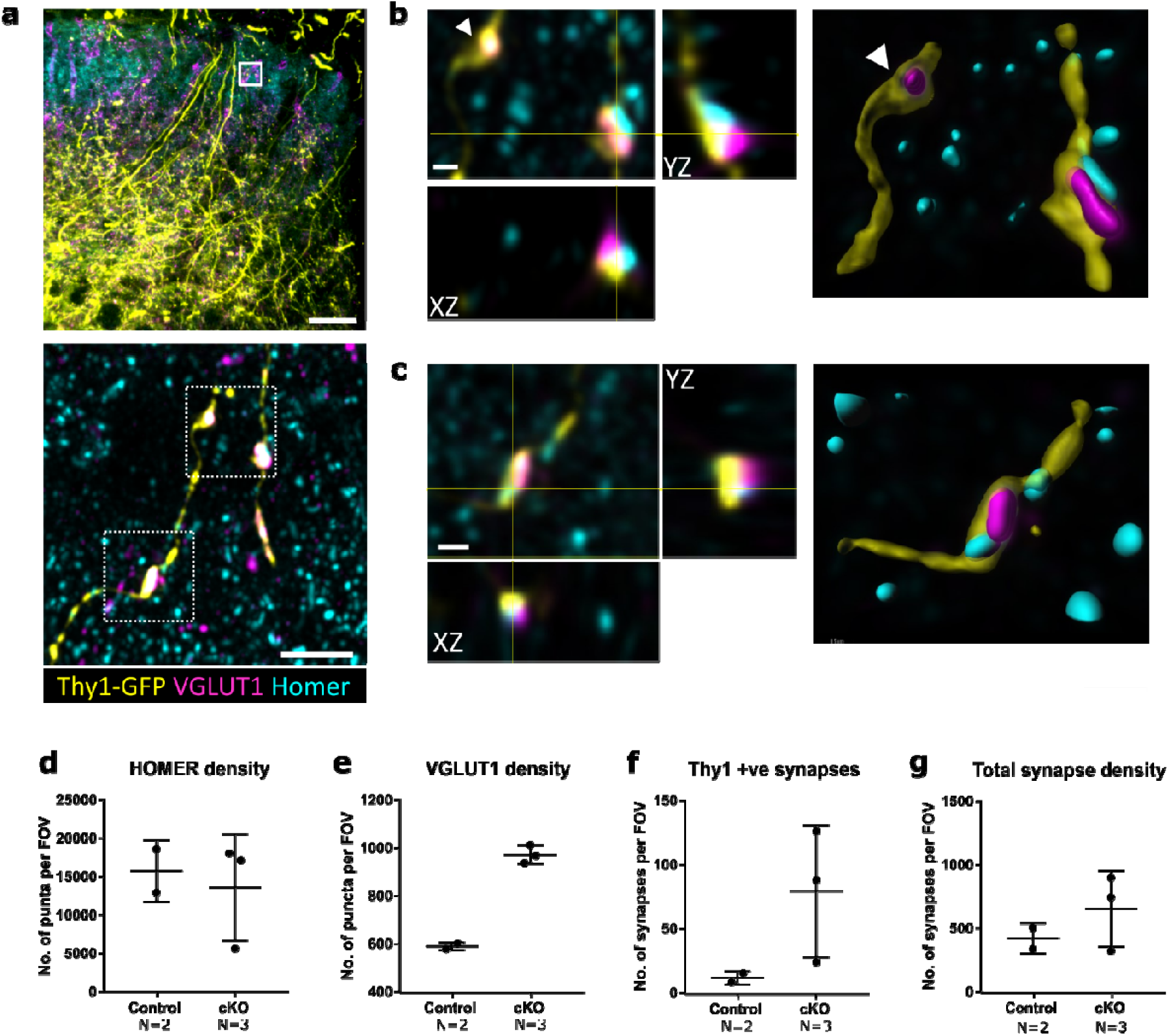
Superficial A-fibres form synapses with pre- and post-synaptic densities. **a**. TOP: Low magnification image, with white box indicating the location of bottom panel image. Scale bar = 50_μ_m. Bottom: Representative maximum projection of super-resolution image from the superficial dorsal horn laminae showing colocalization of Thy1-GFP labelled A-fibre (yellow) with presynaptic VGLUT1 (magenta) and postsynaptic HOMER (cyan). Scale bar = 5_μ_m. **b, c**. Super-resolution images of boxed areas in **a** (bottom panel) with xy, xz, and yz views showing colocalization of Thy1-GFP labelled A-fibre (yellow), presynaptic VGLUT1 (magenta), and postsynaptic HOMER (cyan), as well as surface rendered view on the right. White arrowhead in **b** indicate a Thy1-GFP and VGLUT1 positive presynaptic bouton without postsynaptic HOMER. Scale bars = 1_μ_m. **d**. Homer density does not seem affected in *Tmem16f* cKO animals. **e**. VGLUT1 density seems higher in *Tmem16f* cKO animals, consistent with results in Fig. 3. **f**. Thy1-positive synapse density (as defined by colocalization of Thy1-GFP, VGLUT1, and HOMER) seems to increase in *Tmem16f* cKO animals. **g**. Total synapse number (as defined by colocalization of VGLUT1 and HOMER puncta) seems unchanged in *Tmem16f* cKO animals. FOV= 94_μ_m x 94_μ_m for **d-g**. N-numbers as indicated.

**Figure 3—figure supplement 1.**
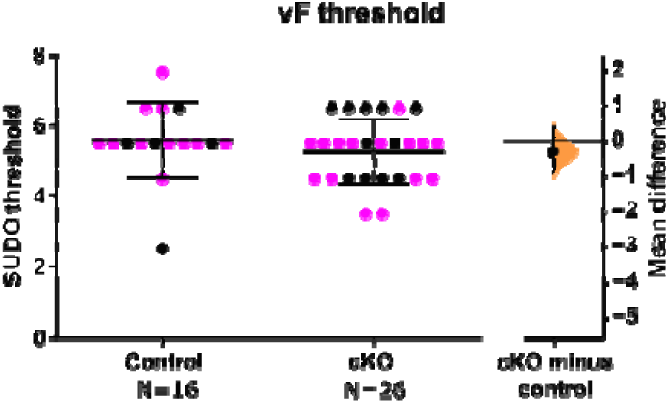
*Tmem16f* cKO does not change vF reflex threshold. Unpaired mean difference −0.29 [95% CI −0.85, 0.39]. N-numbers as indicated. Black and magenta data points indicate females and males respectively.

**Figure 4—figure supplement 1.**
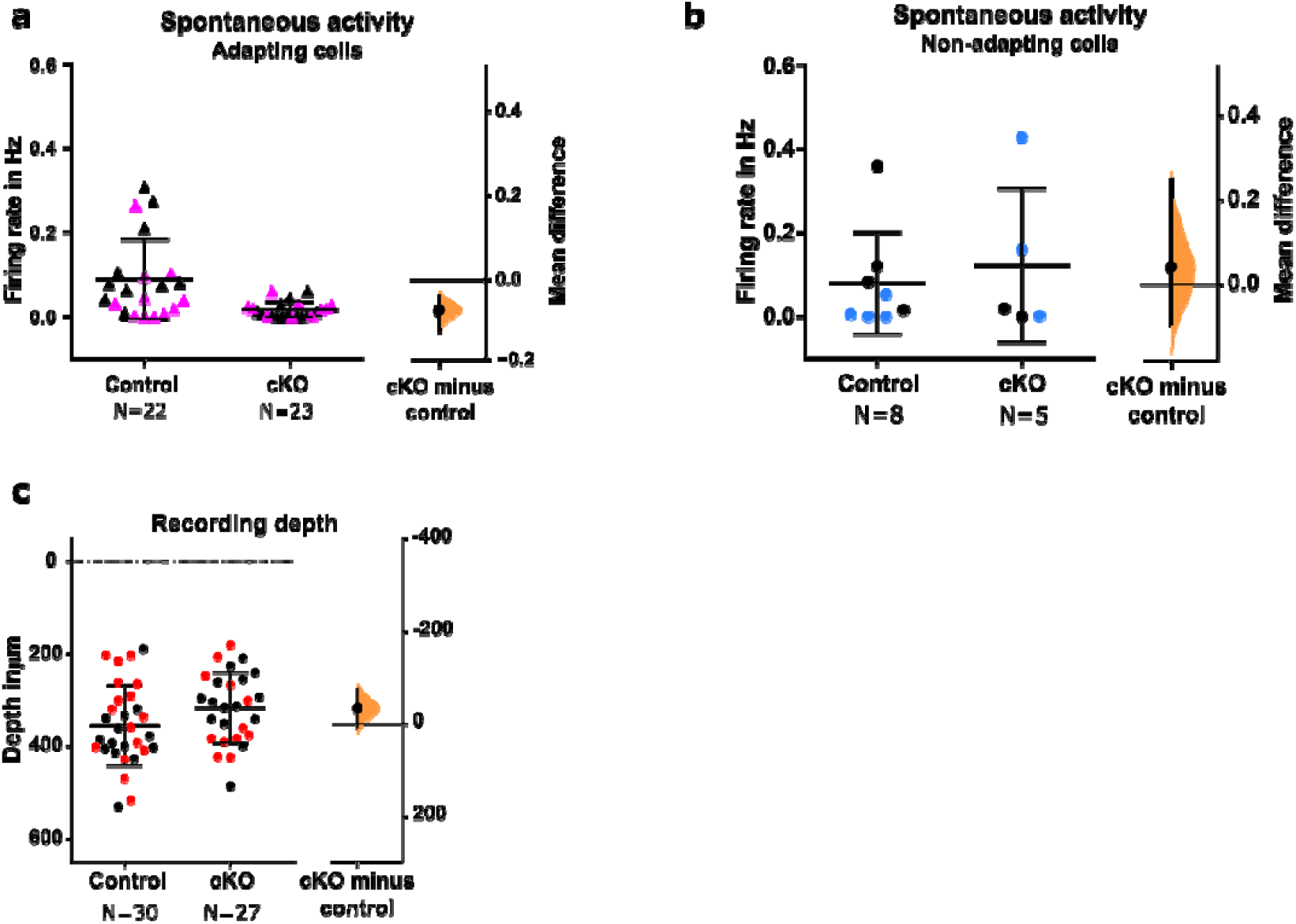
Recording depth and spontaneous activity in control and *Tmem16f* cKO animals. **a** Spontaneous activity decreased in adapting WDR neurons from *Tmem16f* cKO animals, unpaired mean difference −0.072 [95% CI −0.121, −0.039]. **b**. Spontaneous activity was unchanged in non-adapting WDR neurons from *Tmem16f* cKO animals, unpaired mean difference 0.042 [95.00% CI −0.092, 0.25]. **c**. Recording depth of all neurons in control and *Tmem16f* cKO animals, unpaired mean difference - 37.24 [95.00% CI −78.02, 5.73]. N-numbers as indicated. Black and coloured data points indicate females and males.

